# The resistome of common human pathogens

**DOI:** 10.1101/140194

**Authors:** Christian Munck, Mostafa M. Hashim Ellabaan, Michael Schantz Klausen, Morten O.A. Sommer

## Abstract

Genes capable of conferring resistance to clinically used antibiotics have been found in many different natural environments. However, a concise overview of the resistance genes found in common human bacterial pathogens is lacking, which complicates risk ranking of environmental reservoirs. Here, we present an analysis of potential antibiotic resistance genes in the 17 most common bacterial pathogens isolated from humans. We analyzed more than 20,000 bacterial genomes and defined a clinical resistome as the set of resistance genes found across these genomes. Using this database, we uncovered the co-occurrence frequencies of the resistance gene clusters within each species enabling identification of co-dissemination and co-selection patterns. The resistance genes identified in this study represent the subset of the environmental resistome that is clinically relevant and the dataset and approach provides a baseline for further investigations into the abundance of clinically relevant resistance genes across different environments. To facilitate an easy overview the data is presented at the species level at www.resistome.biosustain.dtu.dk.

## Introduction

Over the past decade substantial efforts have been devoted to characterize the antibiotic resistome in different environments(1, 2). Genes conferring resistance to most antibiotics have been found in most environments(1), including pristine environments such as permafrost sediments and isolated cave environments(3, 4). In addition, several clinically relevant resistance genes have recently emerged in human pathogens highlighting the flow of resistance genes from natural environments to human pathogens(5). Yet, an increasing number of studies suggest that substantial barriers to gene transfer across environmental niches exist, implying that genes capable of conferring antibiotic resistance are not easily transferred to human pathogens, especially if selection is absent (6-8). In order to assess the overlap between an environmental resistome and clinically relevant resistance genes a general and comprehensive overview of resistance gene prevalence across human pathogenic species is required. Currently, such overview is only available for subsets of resistance mechanisms or species and have relied on extensive literature mining (9-13). Yet, with the advance of affordable whole-genome sequencing, multiple large-scale sequencing studies of key human pathogens have been conducted(14,15). The data generated in such studies give a detailed insight into the evolution and dissemination of pathogenic strains and enable researchers to develop species-specific genome-based predictions of the resistance genes underlying the resistance phenotypes(16-21). Still, an unbiased analysis of the resistome of pathogenic human isolates is lacking.

In order to get a more complete overview of the antibiotic resistance gene distribution in important human pathogens, we have analyzed all available genomes obtained from human isolates of species commonly known to cause infections. Our study includes genomes available for all the ESKAPE pathogens(22) (*Enterococcus faecium, Staphylococcus aureus, Klebsiella pneumoniae, Acinetobacter baumannii, Pseudomonas aeruginosa* and *Enterobacter spp.)* as well as *Escherichia coli, Mycobacterium tuberculosis, Neisseria gonorrhoeae, Neisseria meningitidis, Streptococus pneumoniae, Streptococcus pyogenes, Enterococcus faecalis, Salmonella enterica, Shigella flexneri* and *Campylobacter jejuni*. These species represent the most common human bacterial pathogens, and antibiotic resistance within many of them is increasing (23).

Based on our genomic analysis we defined a set of clinically relevant resistance genes found in these species. Our analysis provides a comprehensive overview of resistance genes across different pathogens and we provide an easy accessible presentation of the data at www.resistome.biosustain.dtu.dk. This subset of resistance genes will make it easier to identify clinically relevant resistance genes in environmental sequence data and uncover connections between different environmental resistance gene reservoirs and the clinic.

## Materials and methods

### Genome database

All available assembled refseq genomes for the investigated species were downloaded from the National Center for Biotechnology Information (NCBI) (www.ncbi.nlm.nih.gov/refseq/) (downloaded July 2016). Bacterial isolates from humans were identified by searching for the term “human” or “homo” in the host field of the genbank file.

### Identifying resistance genes in the human isolates

Using BLAST, we searched the downloaded genomes for the presence of each gene from the Resfinder resistance gene database (downloaded July 2016) (24), In the BLAST command the following parameters were used: -perc_identity 95 - max_target_seqs 500000000 -taskmegablast-outfmt ′6 std sstrand qlen slen sseq′. The output file was filtered to only include hits with query coverage of >= 90%, queiy coverage was calculated as length/qlen. An overview of the data analysis is given in Supplementary figure 3, at www.resistome.biosustain.dtu.dk.

### Clustering of hits

In order to present the results in a more concise format, the hits were binned into clusters. The resistance gene clusters were based on a clustering of the resistance gene database. The database was clustered using CDhit (80 % identity, 80 % coverage)(39). Using the cluster information from the resistance gene database, the BLAST hits were binned into resistance gene clusters. Each resistance gene cluster was represented by the most common sequence.

### Systematic annotation of the resistance gene clusters

In order to systematically annotate the resistance gene clusters with mechanism of resistance we used the hmm database from ResFam(29). To ensure reliable annotation we first predicted the open reading frame (ORF) in the representative sequence using GeneMarks.hmm(40). Subsequently the mechanism of resistance for the representative protein sequence was annotated with HMMER (hmmscan) using the ResFam database (Resfams-full.hmm)(29, 41).

### Calculation of resistance cluster frequency

For each bacterial species, the frequency of each resistance gene or resistance gene cluster was calculated. This resulted in a table of cluster abundances per species. This table was manually curated to remove genes belonging to the core genome. Genes with high abundance (>93 %) that could be identified as endogenous based on a literature search were removed (see supplementary table 3 for the complete cluster list along with references for removing). The resulting table represents the resistance genes found in common human pathogens referred to as the clinical resistome.

### Generation of the un-clustered database

As the query database contained many highly similar sequences, e.g. single nucleotide variants of the same resistance gene, there were often multiple hits to the same subject. In these cases, the bitscore was used to identify the best hit In addition, as the query database contained highly similar genes with different lengths, e.g. same resistance gene but with an alternative start codon, it was possible for a subject region to have two high scoring hits. To overcome overestimation of resistance gene abundances associated with these variations, hits were grouped according to their location on the target sequence, such that if two hits had start and stop positions within 20 bp. of each other, respectively, only the longest hit was considered.

### Data presentation

All data analysis was done in R, using the packages ggplot2 and vegan(42,43). For the NMDS analysis, Bray-Curtis distances were calculated with binary = TRUE.

For the co-occurrence analysis only resistance gene clusters that had >5% abundance were included. For each pair of resistance gene clusters both frequencies of co-occurrence were calculated, i.e. for clusters A and B, both the frequencies of A+B/A and A+B/B were calculated.

### Flanking region analysis

To confirm genetic linked dissemination, BLAST was used to identify the genomic location of the resistance gene using the subject sequence (sseq) as query. Subsequently, the flanking regions 2 kb up- and downstream from the gene were extracted. The regions were clustered using CD-hit (90 % identity, 90 % coverage). Next, the ORFs in the extracted regions were identified using Genemarks.hmm and clustered into gene families CD-hit (95 % identity, 95 % coverage) and annotated manually. Only the most common region signature is shown in figure 4.

### Data availability

A complete overview of all resistance genes at both single gene (un-clustered) and cluster level for each species is available at the web site www.resistome.biosustain.dtu.dk. In addition, all supplementary information, including fasta files with all un-clustered resistance genes, is available from the same web site.

## Results

We downloaded all 31,073 genomes available for the ESKAPE pathogens plus *E. coli, M. tuberculosis, N. gonorrhoeae, N. meningitides, S. pneumoniae, S. pyrogenes, E.faecalis, S. enterica, S. flexneri* and *C. jejuni* (table 1). From the host descriptor in the genbank files we identified 20,757 (67%) genomes that derived from human isolates. Using BLAST we mined the genomes for resistance genes with the highly curated Resfinder database(24) as the query sequences (Materials and methods).

With this approach we identified 122,463 putative resistance genes across the genomes. Many of these genes represented different variants of the same gene such as the different *tern* or *oxa* betalactamase genes. In order to simplify the results we clustered the resistance genes at 80 % identity (complete unclustered dataset with gene variant resolution is available on www.resistome.biosustain.dtu.dk). We named the resistance gene clusters with a cluster id followed by a common name for the genes in the cluster, e.g. c205-TEM is cluster 205, which contains the *tem* beta-lactamase genes.

In total, 197 unique resistance gene clusters were identified. Of these, some were identified as being native in at least one of the species, e.g. the cluster containing the *oxa-51* beta-lactamase is naturally occurring in *A. baumannii*(25). In order to exclude such native resistance genes we first identified gene clusters with high species abundance (>93% of isolates of a given species) and subsequently mined the literature for evidence that these gene clusters represented native genes (Supplementary table 3). In total 16 gene clusters were removed even though they may contribute to resistance. Importantly, they were only removed from the species they were native to.

After removing the native resistance gene clusters 187 unique resistance gene clusters remained (table 1 and supplementary table 3 www.resistome.biosustain.dtu.dk). These clusters represent 38 % of the 492 resistance gene clusters in the ResFinder database, highlighting that the majority of the resistance genes clusters in the database are not found in sequenced genomes of common human pathogens.

**Table 1.**
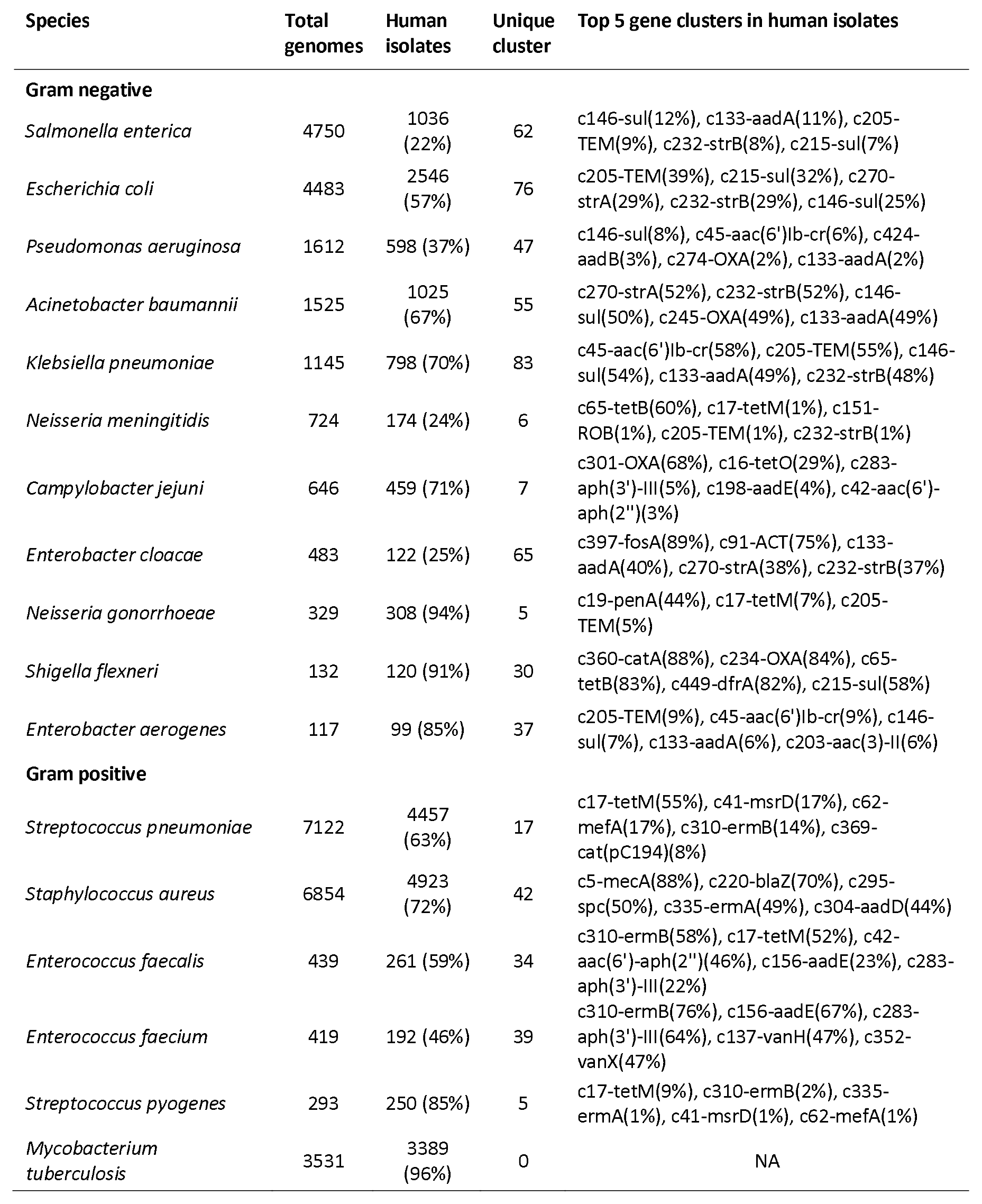
The five most abundant resistance gene clusters in the 17 human pathogens along with general statistics on the species. The total genomes column denotes to total number of refseq genomes for each species. The human isolates column denotes the number of genomes annotated as being from a human host The unique cluster column denotes the number of unique resistance gene clusters found for each species. Only *M. tuberculosis* has no acquired resistance genes. For complete overview visit www.resistome.biosustain.dtu.dk.

Only *M. tuberculosis* did not have any putative resistance genes, which is to be expected, as antibiotic resistance in *M. tuberculosis* is achieved through mutations in native genes as opposed to acquisition of resistance genes(26). The remaining pathogenic species investigated had between 5–62 gene clusters (Table 1). For Gram-negative species the beta-lactamase resistance gene cluster c205-TEM and the sulfonamide resistance gene clusters c146-sul and c215-sul are the most abundant resistance gene clusters. In *E. coli* for instance, 39 % of the genomes were found to carry a beta-lactamase gene belonging to the *tern* family, while c146-sul and c215-sul was found in 25 % and 32 % of the genomes, respectively. Clusters c146-sul and c215-sul contain the*sull* and *sul2* genes, respectively. For the Gram-positive species, the tetracycline resistance cluster c17-tetM and the macrolide resistance gene clusters c335-ermA and c310-ermB are the most abundant resistance gene clusters. For *S. aureus,* 49 % of the genomes contain the c335-ermA cluster and in *S. pneumonia* 55 % contain the c17-tetM cluster and 14 % contain the c310-ermB cluster. Furthermore, 88 % of the *S. aureus* genomes carry the c5-mecA gene cluster responsible for the methicillin resistance Staphylococcus aureus (MRSA) phenotype. *N. meningitidis, N. gonorrhoeae, C. jejuni and S. pyogenes* all have fewer than 10 unique resistance gene clusters suggesting that these species do not harbor many acquired resistance genes compared to other pathogens (Table 1). Comfortingly, these findings are in agreement with prior studies of abundant resistance genes in different bacterial species(9,10,12, 27, 28). Yet, in contrast to previous studies, we employ a consistent analysis pipeline across all available genomes, which can be readily updated with availability of additional genomic information.

In general, there is a trend that resistance gene clusters with many hits are also observed in more species (Figure 1). For instance, the c205-TEM beta-lactamase gene cluster is by far the most disseminated resistance gene cluster as well as one of the gene clusters with most hits. This gene cluster contains 155 unique genes and is found in almost 2000 genomes across 12 of the 17 analyzed species (un-clustered prevalence available on: www.resistome.biosustain.dtu.dk). Other highly disseminated resistance gene clusters found in more than 8 species include: the trimethoprim resistance gene cluster c449-dfrĄ the streptomycin resistance gene clusters c270-strA and c232-strB, the sulfamethoxazole resistance gene clusters c146-sul and c215-sul, and the tetracycline resistance gene clusters c17-tetM, c65-tetB and c67-tetA. In contrast, the c5-mecA gene cluster is only found in *S. aureus*; yet, it is the most abundant resistance gene in the whole dataset, being present in more than 4000 genomes reflecting the selective sequencing of MRSA strains. Although the number and the clonality of the sequenced genomes biases this analysis, it does highlight how some resistance genes have become very successful in disseminating to many different species.

**Figure 1.**
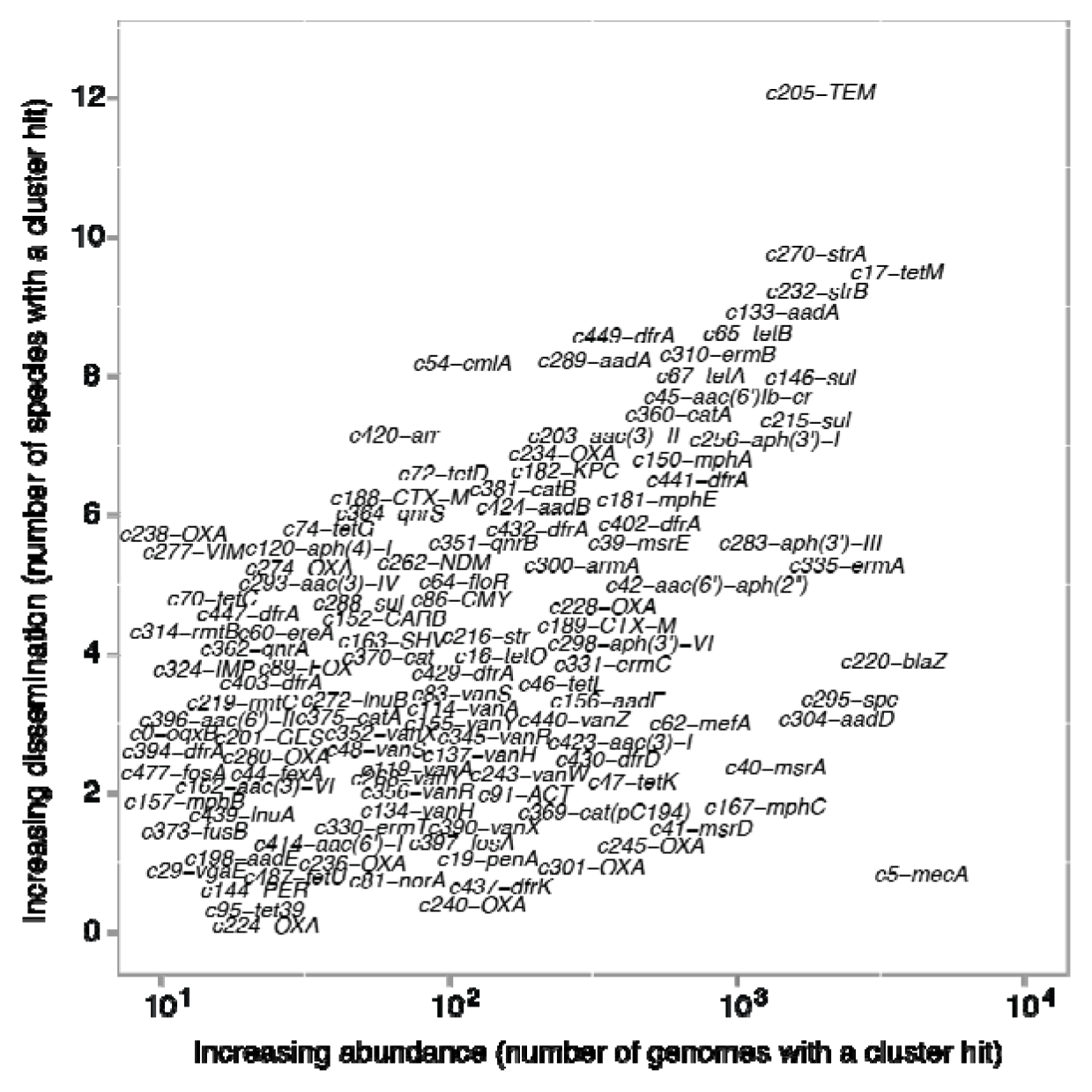
Resistance gene cluster dissemination as a function of abundance. Each of the 187 resistance gene clusters identified in the genomes isolated from humans are plotted according to how many of the 17 analyzed species they were found in and the total number of genomes they were found in. To reduce overplotting, each data point has been jittered on both axes.

## Functional distribution of gene clusters

In order to associate each resistance gene cluster with the antibiotic class that it likely confers resistance to and a mechanism by which it does so, we used ResFam to annotate the resistance gene clusters(29). When stratified by antibiotic class, the abundances of the resistance gene clusters are differentially distributed across the 16 species (*M. tuberculosis* not included). Whereas gene clusters comprising beta-lactam and aminoglycoside resistance genes are found in genomes from most species, gene clusters comprising vancomycin and macrolide resistance genes are only abundant in a small subset of species (Figure 2a). Indeed, vancomycin resistance gene clusters are mainly found in *Enterococcus* spp. and macrolide resistance genes are predominantly found in Gram-positive species. This likely reflects an antibiotic-dependent difference in the dissemination of resistance genes, where the widely used beta-lactam and aminoglycoside antibiotics select for a broad dissemination of beta-lactam and aminoglycoside resistance genes while vancomycin and the macrolide antibiotics are mainly used against *Enterococcus* spp. and *S. aureus* and Gram-positive species, respectively (Figure 2a). Still, macrolide resistance genes do occur in Gram-negative species, which is probably a result of co-selection with other resistance genes.

**Figure 2.**
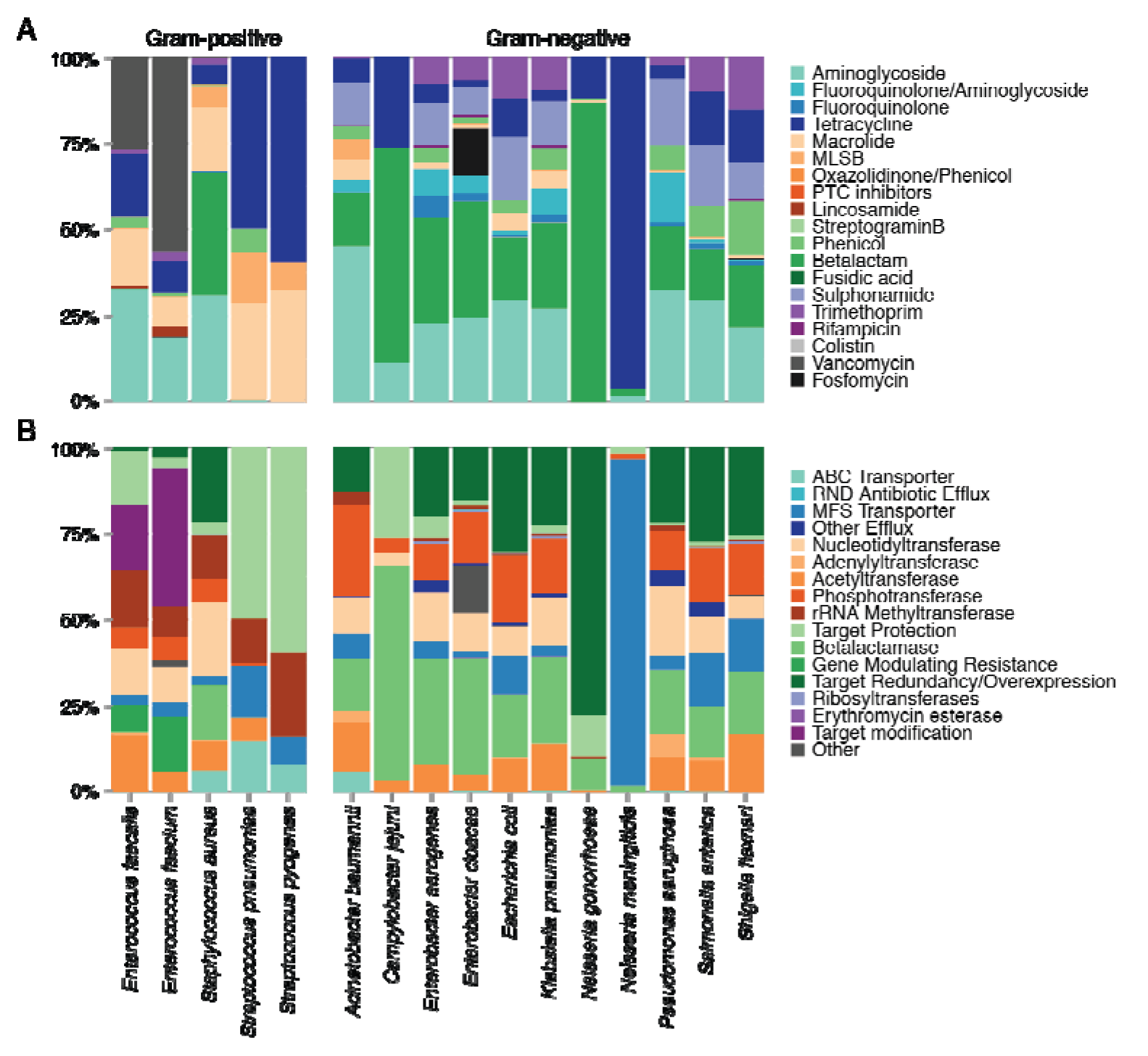
Overview of resistance mechanism and antibiotic category. For all cluster hits across the genomes within a species, the relative distributions of the hits are shown according to the resistance mechanism (A) and the antibiotic category that the cluster confers resistance to (B). The mechanism category is assigned using Resfam and the drug category is implemented from the ResFinder metadata. (Abbreviations: MLSB macrolide, lincosamide, streptogramin B; PTC peptidyl transferase center; ABC ATP-binding cassette; RND resistance nodulation division; MFS major facilitator superfamily)

When stratifying the resistance gene clusters into the different mechanisms underlying the resistance there are also clear divisions between Gram-positive and Gram-negative species. It is interesting to note that resistance gene clusters that encode phosphotransferases, which are commonly involved in aminoglycoside resistance, seem to be more abundant in Gram-negative species than Gram-positive species. Instead, Gram-positive species appear to rely on acetyltranseferases and nucleotidyltransferases to achieve aminoglycoside resistance. Gene clusters encoding rRNA methyltransferases and protection proteins, commonly conferring resistance to peptidyl transferase inhibitors and tetracycline, respectively, are more abundant in Gram-positive species (Figure 2b). For the rRNA methyltransferases, this likely reflects the fact that that the resistance conferred by this mechanism is to antibiotics such as macrolides and streptogramins, both of which are mainly used to treat Gram-positive infections. In the case of the protection proteins the effect of the mechanism is different. This mechanism mainly gives resistance to tetracycline antibiotics, which are effective against both Gram-positive and Gram-negative species. However, while Gram-positive species achieve resistance via ribosomal protection proteins, Gram-negative species generally rely on efflux pumps to clear the drug from the cell(9). This difference might be due to the different cell physiology disfavoring efflux pumps in Gram-positive species. An exception to this division of mechanism between Gram-positive and Gram-negative species is the Gram-negative species *C. jejuni* that commonly has the tetracycline resistance gene *tetO* encoding a ribosomal protection protein.

## Genome level cluster profiles

In order to systematically analyze the distribution of resistance gene cluster hits across different species, we used non-metric multi-dimensional scaling (NMDS) analysis to identify genomes with similar cluster hit profiles (Figure 3). The analysis clearly reveals that Gram-positive and Gram-negative species generally have distinct cluster hit profiles, highlighting that the resistome is largely nonoverlapping between Gram-positive and Gram-negative species. The different cluster hit profiles are most likely driven by multiple factors including differences in cell physiology, antibiotic usage and ability to receive and maintain mobile genetic elements.

**Figure 3.**
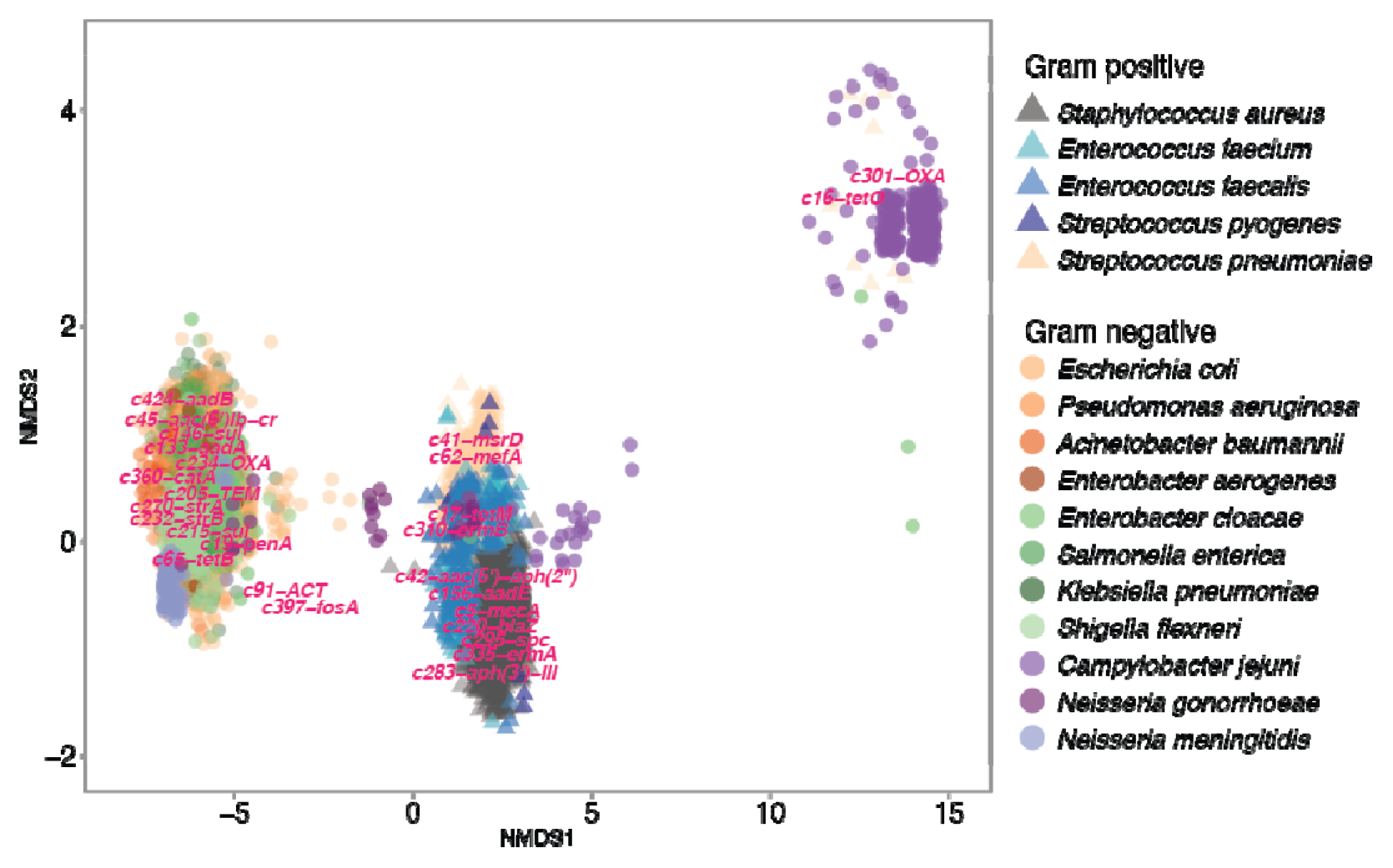
NMDS of the genomes based on the resistance gene profile. NMDS analysis based in the presence or absence of the resistance gene clusters in the individual genomes. The genome observations are overlaid with the cluster scores of the three most abundant clusters for each species. The plot is based on binary Bray-Curtis distances, stress = 0.042. To reduce over-plotting, each data point is jittered on both axes.

By overlaying the most common resistance gene clusters, the analysis also highlights associations between gene cluster and species. For instance, 5. *aureus* genomes are highly clustered by the c5-mecA and c220-blaZ gene clusters (88% and 70 % abundance, respectively), the former causing the MRSA phenotype and only found in *S. aureus.* The *Enterococcus faecalis* genomes cluster by the c310-ermB and c17-tetM gene clusters (58 % and 52 % abundance, respectively), while *E.faecium* is spread out between the c310-ermB and cl56-aadE gene clusters (76% and 67 % abundance, respectively). *Streptococcus pneumoniae* forms two clusters, one driven by the c310-ermB and c17-tetM gene clusters (14 % and 55 % abundance, respectively) overlapping with *E. faecalis*, and one driven by the c41-msrD and c62-mefA gene clusters (both 17 % abundance). In contrast, the cluster hit profiles of Gram-negative genomes are more tightly clustered around the highly disseminated c205-TEM, c146-sul, c215-sul, c270-strA and c232-strB clusters, suggesting a larger extent of resistance gene dissemination within Gram-negative species compared to Gram-positive. Only the *N. meningitides* and a few of the *S. enterica* and *E. cloacae* genomes cluster closely together, the remaining genomes make up one dense cluster with multiple species. This reflects that a major overlap exists between the most common resistance gene clusters across the Gram-negative species, which is also reflected in Table 1 and Supplementary figure 1. Interestingly, *Campylobacter jejuni* species constitute two clusters distinct from the other clusters. This distinction is mainly driven by the c301-OXA gene cluster (68 % abundance), which is unique to *C. jejuni.* In addition, the gene cluster cl6-tetO (29 % abundance) is also a distinct hallmark of *C. jejuni*. Interestingly, the gene clusters cl6-tetO, c283-aph(3’)-III and cl98-aadE (29%, 5% and 4 % abundance respectively), commonly found in *C. jejuni*, are generally associated with Gram-positive genomes and therefore make some *C. jejuni* genomes cluster with the Gram-positive species(30, 31). Notably, the ability of *C. jejuni* to maintain predominantly Gram-positive resistance genes may enable this species to be a bridging species that can link the Gram+ and Gram-resistome. Likewise, for *N. gonorrhoeae*, the predominantly Gram-positive c17-tetM gene cluster (7 % abundance) drives the clustering towards the Gram-positive profiles(31).

## Co-occurrence of resistance gene clusters

As our analyses are performed at the genome level, we were able to identify resistance genes that repeatedly co-occur in the genome of different strains. Genes with strong co-occurrence could be genetically linked and disseminated by a specific integron, transposon or plasmid. For 14 of the 17 species, we generated co-occurrence matrices based on the pairwise co-occurrence of the resistance gene clusters, the remaining three species did not have enough cluster hits to be included in the analysis. We calculated the frequency of co-occurrence as the number of pairwise co-occurrences relative to the total number of occurrences of each gene. To reduce spurious findings, we limited the analysis to genes that were found in more than 5 % of the genomes within a given species. The resulting co-occurrence matrix gives information on the frequency by which a given gene co-occurs with another gene (Supplementary figure 2, see www.resistome.biosustain.dtu.dk). If two genes are always found together, they have complete linkage. If a gene is observed together with another gene in 50 % of the cases, it displays partial linkage and finally, when two genes are never observed together, they display no linkage. It should be noted that, for a gene-pair, A and B, it is possible for gene A to be completely linked to gene B, while gene B is only partially linked to gene A. This would happen if gene B were found in multiple genetically different contexts while gene A was only found in one genetic context, i.e. a specific transposon. While a strong linkage between two or more genes might indicate linked dissemination, it could also result from sequencing of clonal lineages and thus simply reflect the dissemination of a strain rather than a genetic element.

The extent of co-occurrences varies greatly from species to species. *E. coli, A. baumannii, K. pneumoniae, S. enterica, S. flexneri, E. cloacae, E. faecalis, E.faecium* and *S. aureus* have a large co-occurrence network, while *C. jejuni, E. aerogenes, N. gonorrhoeae, P. aeruginosa* and *S. pneumoniae* have a small co-occurrence network with just a few genes (Supplementary figure 2, see www.resistome.biosustain.dtu.dk). *S. pyogenes, N. meningitidis* and *M. tuberculosis* do not have enough gene clusters above the threshold and therefore the co-occurrence network could not be computed.

For some gene clusters, the co-occurrence is well understood e.g. the cooccurrence of *strA* and *strB*, which combined give high level streptomycin resistance(32). Similarly, the different *van* gene clusters conferring vancomycin resistance in enterococci are strongly co-occurring, as resistance is achieved through the concerted action of these genes(10).

The co-occurrence analysis also recapitulates known co-dissemination patterns driven by linkage on mobile genetic elements. For instance, the macrolide resistance gene cluster c335-ermA in *S. aureus* is located just upstream of the spectinomycin resistance gene cluster c295-spc (Figure 4A). The two genes are a part of transposon Tn554, which in turn is a part of a larger pathogenicity island including the c5-mecA and the c304-aadD clusters(33, 34). As a result the clusters c295-spc, c335-ermA and c304-aadD are all strongly linked to each other as well as to c5-mecA; however, c5-mecA is found in many other contexts and therefore not strongly linked to the three other resistance genes (Figure 4A).

In *E. coli* there is a strong linkage between the streptomycin/spectinomycin resistance gene cluster c289-aadA and the trimethoprim resistance cluster c402-dfrA, and a further linkage to the sulfamethoxazole resistance gene cluster c146-sul. These resistance genes are commonly found together in class 1 integrons in strains of both human and animal origin (35) (Figure 4B). Interestingly, of the 284 genomes where clusters c289-aadA and c402-dfrA co-occur, 242 (85 %) are found in an integron with an IS26 insertion truncating the *intl*1 gene (Figure 4B).

**Figure 4.**
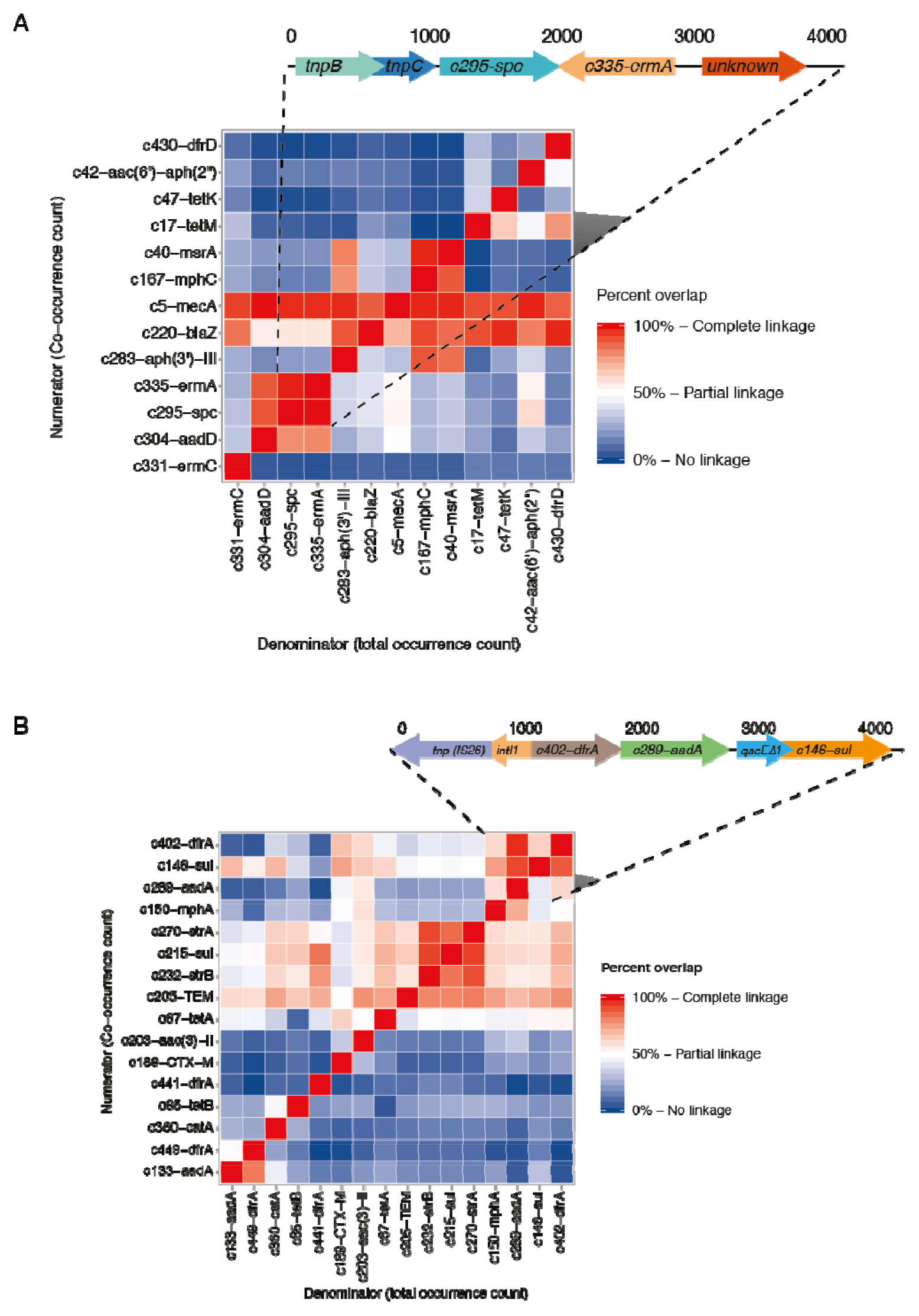
Co-occurrence matrix. A) Co-occurrence matrix for S. aureus, the macrolide resistance gene c335-ermA is found in 2393 (49 %) of the *S. aureus* genomes and in 2382 (99.5 %) of the cases it is found together with the c295-spc gene, most commonly in the configuration depicted in the figure as part of the transposon Tn554, where the two genes are located next to each other. Co-occurrences are only calculated for genes present in minimum 5 % of the analyzed genomes. B) Co-occurrence matrix for *E. coli* the aminoglycoside resistance gene cluster c289-*aadA* is found in 284 (12 %) of the *E. coli* genomes and in 242 (85 %) of the cases it is found together with the c402-*dfrA* gene cluster, most commonly in the configuration depicted in the figure as a class 1 integron with an IS26 insertion truncating the *intll* gene. Co-occurrences are only calculated for genes present in minimum 5 % of the analyzed genomes.

In addition to identifying linked resistance genes, the co-occurrence analysis also identifies genes that are rarely or never found together. This can be caused by resistance genes being mobilized on different plasmids belonging to the same incompatibility group or by a high fitness cost of simultaneous carriage of two genes. For instance, hits to cluster c65-tetB are not found to co-occur with hits to cluster c67-tetA. A possible explanation for this observation could be a lack of selective benefit from simultaneous carriage of *tetA* and *tetB* due to their identical phenotype. The same phenomenon is observed for the three *dfrA* clusters c402, c441 and c449 (Figure 4B).

## Discussion

The declining cost of whole-genome sequencing enables detailed studies of the evolution and dissemination of pathogenic bacteria. Here, we present the results of a genome-wide identification of resistance genes in publicly available genomes of key pathogens isolated from humans. We have identified close homologues of the main genetic determinants of antibiotic resistance in 17 common human pathogens. We found that 38% of the resistance gene clusters from the query database were found in bacterial pathogens isolated from humans, highlighting that most resistance gene clusters were not present in the sequenced genomes.

As our analysis is done *in silico* with no biochemical validation we used stringent thresholds requiring a hit to have a minimum of 95% sequence identity and a minimum of 90% coverage to a query. This ensures, that hits have a high likelihood of being true resistance genes. However, although our analysis identifies close homologues of resistance genes it does not necessarily imply phenotypic resistance, as genes may not always be expressed.

To further ensure a reliable prediction of resistance genes we exclusively focused on non-mutational resistance, although we acknowledge that clinically important resistances also occur via mutations in native genes. In addition, some species may be intrinsically resistant due to native genes conferring resistance, and while this is an important phenomenon, such intrinsic resistance mechanisms are not included in our study.

Importantly, our analysis does not represent a study of horizontal gene transfer. We used the ResFinder database as a reference database for resistance genes as this is a manually curated and actively maintained database of acquired resistance genes(24). Other studies have focused on large-scale identification of HGT typically requiring computational intensive methods (6, 36).

In addition to the Resfinder database other databases of resistance genes, with varying degrees of curation and maintenance, exist(37). Currently, the Comprehensive Antibiotic Resistance Database (CARD) is the largest database including resistance-conferring variants of native genes. However, when comparing the CARD subset of acquired resistance genes to the ResFinder database, we found that CARD contained many native genes, for instance 12% of the genes that were unique to CARD compared to ResFinder (representing 43 gene clusters) were native to *E. coli* K12, which is broadly considered to be a sensitive organism. Such genes would lead to overestimation of resistance gene abundance in our analysis (see Supplementary table 4 for a complete list of CARD exclusive genes). In contrast, just 16 clusters from the ResFinder database were identified as native resistance genes.

Currently, many of the available genomes represent sequencing efforts directed towards specific phenotypes such as MRSA or carbapenem-resistant *K. pneumonia*. While these highly resistant pathogens are a major threat to human health, their overrepresentation in publicly available databases limit studies into the general trends of antibiotic resistance evolution and spread. To obtain a more unbiased view of the global emergence and spread of resistance genes, whole genome sequences from organisms isolated as a part of routine diagnostics in medical institutions around the world should be deposited to public databases. Alternatively, sequencing of key pathogens, representatively sampled at regular intervals, could provide valuable data on the trends of resistance gene evolution and dissemination.

As studies have found that resistance genes can be found in all environments it is relevant to identify the genes that emerge in common human pathogens(1). With our generalized analysis, we have generated a subset of resistance genes found in common pathogens of human origin. This database represents the current subset of resistance genes that are particular relevant to human health. We believe that the database will be useful in assessing the interaction between environmental and clinical resistomes based on metagenomic and genome sequencing studies. Further updates of the database will identify the movement of genes from the general resistome into the clinically relevant resistome. In turn, this can be used to rank the risk-potential of different reservoirs of antibiotic resistance genes(38). Furthermore, if representative collections of bacterial genomes from clinical microbiological departments around the globe were available for analysis, approaches such as the one presented here could be used to more effectively monitor the changing patterns of antibiotic resistance genes and identify the genetics determinants that contribute most to the dissemination of antibiotic resistance.

## Availability

Full resistance gene analysis along with supplementary data are available at www.resistome.biosustain.dtu.dk.

## Acknowledgements

This research was funded by the EU H2020 ERC-20104-STG LimitMDR (638902) and the Danish Council for Independent Research Sapere Aude programme DFF - 4004-00213. MOAS acknowledges additional funding from the Novo Nordisk Foundation and The Lundbeck Foundation.

